# Single-Cell Spatial Proteomics Analyses of Head and Neck Squamous Cell Carcinoma Reveal Tumor Heterogeneity and Immune Architectures Associated with Clinical Outcome

**DOI:** 10.1101/2021.03.10.434649

**Authors:** Katie E. Blise, Shamilene Sivagnanam, Grace L. Banik, Lisa M. Coussens, Jeremy Goecks

## Abstract

There is increasing evidence that the spatial organization of cells within the tumor-immune microenvironment (TiME) of solid tumors influences survival and response to therapy in numerous cancer types. Here, we report results and demonstrate the applicability of quantitative single-cell spatial proteomics analyses in the TiME of primary and recurrent human papillomavirus (HPV)-negative head and neck squamous cell carcinoma (HNSCC) tumors. Single-cell compositions of a nine patient, primary and recurrent (n=18), HNSCC cohort is presented, followed by deeper investigation into the spatial architecture of the TiME and its relationship with clinical variables and progression free survival (PFS). Multiple spatial algorithms were used to quantify the spatial landscapes of immune cells within TiMEs and demonstrate that neoplastic tumor-immune cell spatial compartmentalization, rather than mixing, is associated with longer PFS. Mesenchymal (αSMA^+^) cellular neighborhoods describe distinct immune landscapes associated with neoplastic tumor-immune compartmentalization and improved patient outcomes. Results from this investigation are concordant with studies in other tumor types, suggesting that trends in TiME cellular heterogeneity and spatial organization may be shared across cancers and may provide prognostic value in multiple cancer types.

## INTRODUCTION

Tumor microenvironments, comprising both neoplastic tumor cells and recruited stromal cells of various lineages, including a diverse assemblage of immune, mesenchymal, and vascular cells, play a key role in both de novo progression of tumors and regulating response to therapies^1–3^. Numerous studies have reported that, in addition to the types and quantities of cells present in the tumor immune-microenvironment (TiME), the spatial organization of the TiME is prognostic for survival and response to therapy in multiple cancer types^4–13^. Metrics that quantify this spatial organization can range from simple density ratios within specific tumor regions^14^, such as the Immunoscore^15^, a now commonly used biomarker for colorectal tumor staging, to more complex measures that account for the precise locations of specific cells relative to other cells, such as mixing scores^4^ and cellular neighborhood measures^7^. These more advanced spatial quantifications are a result of emerging single-cell multiplex tissue imaging modalities^4, 6, 7, 16–18^, which provide detailed phenotypic and effector proteomic markers for each cell, while maintaining the spatial architecture of the tissue assayed. Knowing the precise locations of cells in the TiME enables a deeper understanding of how cells interact within the tumor, as both direct and indirect cell signaling mechanisms require cells to be near, if not directly adjacent to one another^19^. This understanding can aid treatment decisions, as many therapies require spatial proximity of specific cell types for efficacy^10^. Given that single-cell imaging technologies are still relatively new, there is much to be discovered regarding how the spatial organization of cells within the TiME relates to clinical outcome and may be used for patient stratification decisions for therapy.

Head and neck squamous cell carcinoma (HNSCC) is the sixth leading form of cancer worldwide^20^, and it accounts for more than 10,000 deaths per year in the US alone^21^. While patients harboring human papillomavirus (HPV) within neoplastic cells tend to exhibit a better prognosis, their HPV-negative [HPV(-)] counterparts typically exhibit T cell suppressive TiMEs and have a significantly greater risk of recurrence and shorter 3-year survival^22–24^. There is a critical need to improve understanding of HNSCC TiMEs to enable better patient stratification for therapy, as well as identify new targets that could be leveraged for therapeutic intervention to improve outcome, particularly for patients with HPV(-) tumors who currently lack promising therapeutic options. We previously developed a multiplex immunohistochemistry (mIHC) imaging platform to aid studies investigating the immune contexture of solid tumors and their response to therapies at the single-cell level^16, 17^. Using a sequential antibody staining protocol, detection of 12-30 proteins can be enumerated at single-cell resolution across a single formalin-fixed paraffin-embedded (FFPE) tissue section. This enables single-cell phenotyping of discrete leukocyte lineages, and importantly, reveals their spatial relationships with other cells in the tissue section. Utilizing this mIHC approach on a small cohort of eighteen HPV(-) primary and matched recurrent HNSCC tumor samples collected from nine patients, we previously reported immune contextures associated with disease recurrence, most notably that myeloid inflamed profiles in primary tumors exhibited shorter progression free survival (PFS) compared to lymphoid inflamed profiles^16, 17^.

In this study, we have significantly extended our prior analysis of this cohort, focusing on tumor heterogeneity and compositional changes from primary to recurrent tumors, in addition to using multiple spatial algorithms to quantify the spatial organization of the TiMEs. We then correlated these spatial features with PFS and identified TiME architectures that may be important for therapeutic decision making. Overall, we found increased neoplastic tumor-immune cell spatial compartmentalization in primary tumors to be associated with longer PFS. These tumors also contained alpha smooth muscle actin (αSMA^+^) cells with more organized structure located near T and B cells, as well as near cells involved in antigen presentation. Our results are concordant with those from other studies, indicating that the features identified herein are likely shared and prognostic across cancer types.

## RESULTS

One to three regions of 2500^2^ µm^2^ from each of the nine patients primary and matched recurrent tumor resections (n=18) were analyzed, for a total of 47 regions (**Table 1, Figure 1a**). For this study, we utilized a gating strategy with thirteen lineage or functional protein biomarkers to classify cells as neoplastic tumor cells, stromal cells (mesenchymal), or one of seven different leukocyte subtypes spanning lymphoid and myeloid lineages (**Table 2, Supplemental Figure 1a**). We investigated tumor heterogeneity both within and across patient samples, quantified the cellular spatial relationships within the TiME using a mixing score and performed a neighborhood clustering method to describe the association between TiME spatial architecture, clinical features, and PFS.

**Figure 1.**
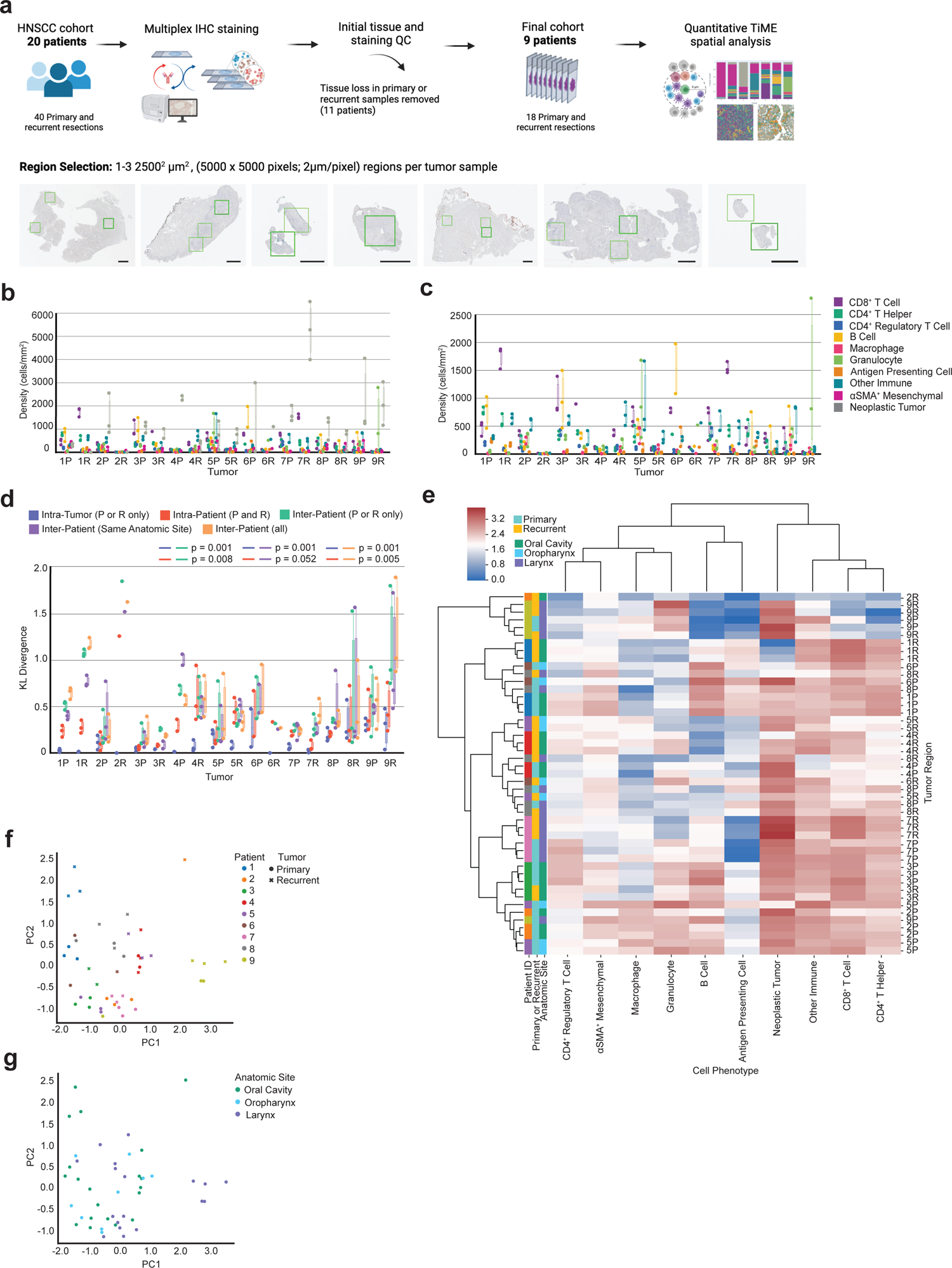
Heterogeneity across patients and tumor regions. **a**, Cohort and tissue region selection overview. One to three regions of 2500^2^ µm^2^ were assayed with mIHC per eighteen tumor resections and are represented by the green boxes in the tissue images. The black line corresponds to 2500 µm for each image. **b**, Density (cells/mm^2^) of each cell type present per individual primary (P) and recurrent (R) tumor. Each dot represents a single tumor region (n=47). **c**, Density (cells/mm^2^) of each immune cell type present per individual primary (P) and recurrent (R) tumor. Each dot represents a single tumor region (n=47). **d**, Box plot of the Kullback-Leibler divergences from a single tumor region’s cellular distribution compared to: the tumor’s average cellular distribution [Intra-Tumor (P or R only)], the patient’s average cellular distribution [Intra-Patient (P and R)], the cohort’s average cellular distribution across tumors of the same timepoint [Inter-Patient (P or R only)], the cohort’s average cellular distribution across tumors of the same anatomic site [Inter-Patient (Same Anatomic Site)], the cohort’s average cellular distribution across all tumors from all patients [Inter-Patient (all)]. P-values calculated using a one-way ANOVA multi-group significance test followed by a Tukey honestly significant difference post-hoc test. **e**, Heatmap of cellular composition across tumor regions. Rows are individual tumor regions that are ordered by the hierarchical clustering of their cellular composition. Columns are the cell types used as clustering features. Composition was normalized using a log10+1 transformation before clustering. Leftmost three columns are color coded by patient, tumor timepoint, and anatomic site. **f**, Principal component analysis on cellular density following a log10+1 transformation. Each point (n=47) represents one tumor region and is colored by patient. The shape of each point denotes primary or recurrent status. **g**, Principal component analysis on cellular density following a log10+1 transformation. Each dot (n=47) represents one tumor region and is colored by the anatomic resection site.

**Table 1.** Patient and tumor characteristics. Adapted from Banik et al^17^ to show de-identified patient characteristics.

| Patient ID | Anatomic Site of Resection | Primary Tumor TNM Stage | Therapy Following Primary Resection | Gender | Race | Alcohol History | Smoking History | HPV Status | Progression Free Survival (days) |
| --- | --- | --- | --- | --- | --- | --- | --- | --- | --- |
| 1 | Oral Cavity | 2 | Cisplatin + Radiation | Male | White | Yes | Yes | Negative | 804 |
| 2 | Oral Cavity | 4 | Cetuximab + Radiation | Male | White | Yes | Yes | Negative | 123 |
| 3 | Oral Cavity | 1 | Cisplatin + Radiation | Female | White | Yes | No | Negative | 1447 |
| 4 | Oral Cavity | 1 | Cisplatin + Radiation | Female | White | No | No | Negative | 246 |
| 5 | Oropharynx | 4 | Cetuximab + Radiation | Female | White | Yes | No | Negative | 202 |
| 6 | Oropharynx | 4 | Cisplatin + Radiation | Female | Asian | No | No | Negative | 188 |
| 7 | Larynx | 4 | Cisplatin + Cetuximab + Radiation | Female | White | Yes | Yes | Negative | 409 |
| 8 | Larynx | 3 | Cisplatin + Radiation | Female | White | Yes | Yes | Negative | 1033 |
| 9 | Larynx | 4 | Cisplatin + Radiation | Female | White | Yes | Yes | Negative | 83 |

**Table 2.** mIHC cell phenotype classification. Antibody panels used to hierarchically phenotype cells into one of ten phenotype classifications. Additional functional markers detected are listed at the bottom.

| Cell Phenotype | Antibody Markers |
| --- | --- |
| CD8<sup>+</sup> T Cell | CD45<sup>+</sup> CD20<sup>-</sup> CD3<sup>+</sup> CD8<sup>+</sup> |
| CD4<sup>+</sup> T Helper | CD45<sup>+</sup> CD20<sup>-</sup> CD3<sup>+</sup> CD8<sup>-</sup> FOXP3<sup>-</sup> |
| CD4<sup>+</sup> Regulatory T Cell | CD45<sup>+</sup> CD20<sup>-</sup> CD3<sup>+</sup> CD8<sup>-</sup> FOXP3<sup>+</sup> |
| B Cell | CD45<sup>+</sup> CD20<sup>+</sup> |
| Macrophage | CD45<sup>+</sup> CD20<sup>-</sup> CD3<sup>-</sup> CD66B<sup>-</sup> CD68<sup>+</sup> |
| Granulocyte | CD45<sup>+</sup> CD20<sup>-</sup> CD3<sup>-</sup> CD66B<sup>+</sup> |
| Antigen Presenting Cell | CD45<sup>+</sup> CD20<sup>-</sup> CD3<sup>-</sup> CD66B<sup>-</sup> CD68<sup>-</sup> MHCII<sup>+</sup> |
| Other Immune | CD45<sup>+</sup> CD20<sup>-</sup> CD3<sup>-</sup> CD66B<sup>-</sup> CD68<sup>-</sup> MHCII<sup>-</sup> CD8<sup>-</sup> FOXP3<sup>-</sup> |
| αSMA<sup>+</sup> Mesenchymal | CD45<sup>-</sup> PANCK<sup>-</sup> αSMA<sup>+</sup> |
| Neoplastic Tumor | CD45<sup>-</sup> PANCK<sup>+</sup> |
| **Additional Functional Markers:** PD-1, PD-L1, Ki-67 |  |

### Single-cell proteomic analyses reveal varying degrees of tumor heterogeneity

To quantify how cellular composition varied across tumor regions, we assessed tumor heterogeneity at multiple levels, including intra-tumoral, intra-patient, and inter-patient cellular heterogeneity by calculating Kullback-Leibler (KL) divergences for each region, performing hierarchical clustering, and conducting a principal component analysis (PCA). The density of each cell type per region was measured for all eighteen tumor specimens by taking the count of each cell type divided by the measured tissue area in mm^2^ (**Figure 1b,c**). We then calculated the coefficient of variation per cell type for each tumor, and averaged these values to quantitatively describe the cell types contributing most to intra-tumoral heterogeneity within the cohort. The coefficient of variation is defined as the standard deviation divided by the mean, and it provides a normalized measure of variability for comparison across cell types with large differences in densities. On average, B cells exhibited the greatest coefficient of variation across the cohort relative to other cell types (**Table 3**). This is likely due to the fact that B cells were frequently observed to be spatially clustered together, resulting in regions of either high B cell density or low B cell density despite being collected from the same tumor (**Supplemental Figure 1b**).

**Table 3.** Coefficient of variation. The coefficient of variation was calculated per tumor per cell type and then averaged across all tumors (n=16). Tumors with only one region (n=2) were removed from this analysis. Cell types are ordered from largest coefficient of variation to smallest coefficient of variation.

| Cell Phenotype | Average Coefficient of Variation |
| --- | --- |
| B Cell | 0.658 |
| Macrophage | 0.563 |
| αSMA<sup>+</sup> Mesenchymal | 0.545 |
| Neoplastic Tumor | 0.541 |
| Granulocyte | 0.523 |
| Antigen Presenting Cell | 0.519 |
| Other Immune | 0.402 |
| CD8<sup>+</sup> T Cell | 0.384 |
| CD4<sup>+</sup> T Helper | 0.376 |
| CD4<sup>+</sup> Regulatory T Cell | 0.311 |

To further quantify and assess tumor heterogeneity both within and across patients, we calculated the KL divergence of each tumor region from five average cell type distributions. KL divergence is a relative measure of how similar two distributions are, with larger values reflecting less similarity between the distributions and smaller values reflecting more similarity between the distributions. This measure has been used previously to quantify tumor heterogeneity^6^. By calculating and comparing the divergences of each tumor region from multiple average cell type distributions, we were able to assess heterogeneity within and across tumors and patients. Overall, we observed that heterogeneity was lower across regions from the same tumor and tumors from the same patient (primary or recurrent), while higher across tumors from different patients. This is evidenced by smaller intra-tumoral and intra-patient KL divergence values for the majority of tumor regions (**Figure 1d**).

The cellular distributions used to calculate the five KL divergence values per tumor region were (1) the average cellular distribution across all regions sampled from the same tumor [“Intra-Tumor (P or R only)”]; (2) the average cellular distribution across the patient’s primary and recurrent tumors [“Intra-Patient (P and R)”]; (3) the average cellular distribution across all tumors in the cohort collected from the same timepoint [“Inter-Patient (P or R only)”]; (4) the average cellular distribution across all tumors in the cohort resected from the same anatomic site [“Inter-Patient (Same Anatomic Site)”], and; (5) the average cellular distribution across all tumors collected from all patients in the cohort, regardless of primary or recurrent status or anatomic site [“Inter-Patient (all)”]. By comparing the relative KL divergence values to each other, we found tumor regions to be more similar to regions sampled from the same tumor and patient than regions collected from tumors of other patients. Notably, we found no significant difference between “Inter-Patient (Same Anatomic Site)” and “Inter-Patient (all),” indicating that tumor regions diverged by the same degree from regions sampled at the same anatomic site as they did from regions sampled at all three anatomic sites (oral cavity, oropharynx, larynx) of the head and neck region. Finally, given the large proportion of neoplastic tumor cells comprising the TiME for many of the tumor regions, we assessed KL divergence using only the distribution of immune cells present and found similar results (**Supplemental Figure 1c**). This indicates that immune cell composition is more similar within regions from the same patient than across regions collected from different patients.

To further investigate intra-patient heterogeneity, we performed unsupervised hierarchical clustering on the 47 tumor regions based on their normalized density composition (**Figure 1e**). We found that two patients (3, green; 7, pink) contained all tumor regions clustering together, independent of primary or recurrent state. These patients also had the smallest intra-patient KL divergence values (**Figure 1d**), indicating that the cell densities of these patients’ primary and recurrent tumors were similar to each other. Three patients (2, orange; 4, red; 9, yellow) contained nearly all regions clustered together. The remaining four patients’ tumors exhibited greater degrees of intra-patient heterogeneity, as demonstrated by the distance between primary and recurrent tumor regions on the clustered heatmap (**Figure 1e**). Overall, we found that regions evaluated from the same patient tended to cluster together more than regions evaluated from different patients (**Figure 1e**), indicating increased heterogeneity between patients as compared to between samples from the same patient. We also examined whether tumor regions clustered by the anatomic resection site and found that the clusters formed did not group by site. These results, in addition to those of our KL divergence analyses, indicate that anatomic site was likely not the main contributor of cellular heterogeneity in this cohort. PCA results also supported these observations (**Figure 1f,g**).

### TiME cellular composition altered by therapy

Multiple studies have reported differences in TiME cellular makeup^25, 26^ and tumor clonal diversity^27^ between primary and recurrent tumors. To assess whether any immune contexture changes occurred following post-operative therapy in our cohort, we analyzed the cellular composition of primary tumors as compared to their recurrent tumors. All patients received a combination therapy of cisplatin and/or cetuximab accompanied by radiation following surgical resections of their primary tumors. We used the average density of each cell type present across regions for a given tumor and compared primary tumor composition to their matched recurrent tumor composition.

While we did not observe any significant differences in cell density between primary and recurrent tumors (p>0.112), we did find that all patients experienced a decrease in the density of B cells from their primary to recurrent tumors (**Figure 2a,b**). This result is supported by a recent study that found that a large cohort of HNSCC patients experienced a decline in B cells from primary to recurrent tumors, and this reduction was dependent upon patients receiving adjuvant chemoradiotherapy^28^. We found no common differences across all patients for any other cell type, although eight of the nine patients saw a decrease in the density of neoplastic tumor cells from their primary to recurrent tumors (**Figure 2b**).

**Figure 2.**
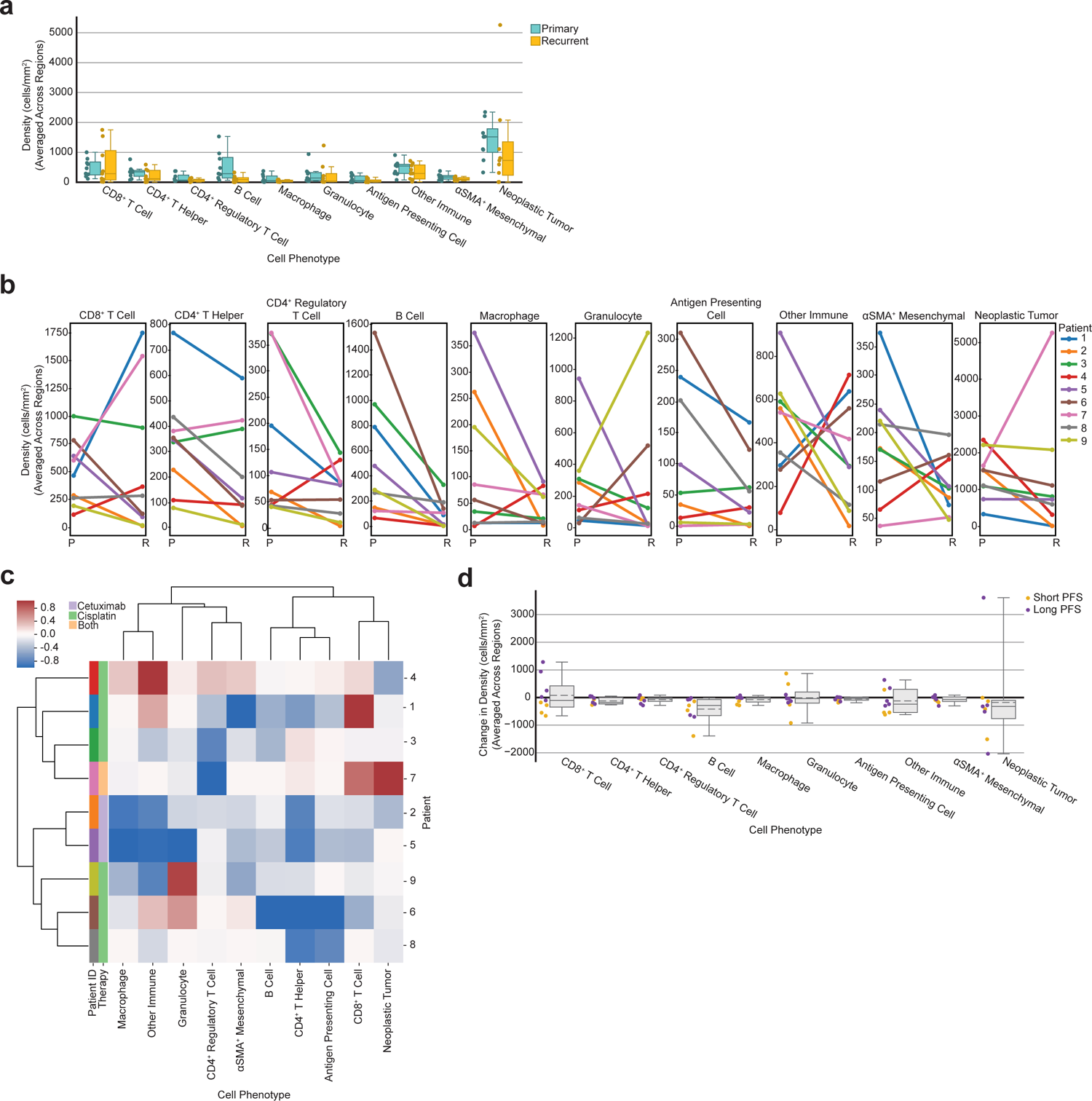
Tumor cellular composition changes following therapy. **a**, Box plot showing the average density of each cell type split by primary and recurrent status. Each dot represents the density of that cell type for one tumor, averaged across regions (n=9 primary tumors, n=9 recurrent tumors). **b**, Line plots showing the change in cellular composition from primary to recurrent tumors for each cell type. Each dot represents the density of that cell type for one tumor, averaged across regions. Lines are colored by patient. **c**, Heatmap of average change in cellular composition from primary to recurrent tumors for each patient. Rows are individual patients that are ordered by the hierarchical clustering of their change in TiME cellular composition (averaged across regions). Columns are the cell types used as clustering features. Compositional change was normalized [-1,1] before clustering (see Methods). Leftmost two columns are color coded by patient followed by therapy administered. **d**, Box plot showing the average change in density of each cell type for each patient (n=9) colored by short-term or long-term progression free survival, splitting on median progression free survival.

We then assessed whether patients exhibited similar compositional differences in primary and recurrent TiMEs by performing unsupervised hierarchical clustering on the normalized average difference in cellular composition for each cell type. This resulted in two groups of patients (**Figure 2c**). Interestingly, the two patients that received cetuximab (2, orange; 5, purple), rather than cisplatin, clustered together within one of these groups and were the only two patients to experience a decrease in the density of every cell type following therapy. Across the cohort these patients had the greatest decrease in the density of macrophages, granulocytes, and other CD45^+^ immune cells present from their primary tumors to their recurrent tumors following therapy (**Figure 2b,c**). Interestingly, the one patient that received both cisplatin and cetuximab (7, pink) was present in the other cluster from the two patients that received only cetuximab. This was the only patient to experience an increase in the density of neoplastic tumor cells (**Figure 2b,c**). This patient also experienced the second largest increase in CD8^+^ T cells as well as the greatest decrease in CD4^+^ regulatory T cells, potentially indicating a pro-inflammatory response to—or despite—increased neoplastic tumor cell density.

Altogether, these results indicate that shared trends in TiME composition changes from primary to recurrent tumors specific to therapy exist, and regardless of therapy, all patients exhibited a significant decrease in B cells from primary to recurrent tumors.

Approximately half (n=4) of the patients in the cohort experienced an increase in CD8^+^ T cell density while the other half (n=5) experienced a decrease in CD8^+^ T cell density following therapy. This was the only cell type that increased in density for nearly half of the cohort and decreased for the other half. To determine if there was a survival advantage for patients that experienced this increase, we split our cohort into short-term or long-term survivor groups using median PFS and observed that all patients who experienced an increase in CD8^+^ T cell density from their primary to recurrent tumors were long-term survivors (**Figure 2d**). Interestingly, the density of CD8^+^ T cells in the primary tumor alone did not associate with PFS (p=0.829). Prior research has revealed that increased CD8^+^ T cell abundance in the primary tumor is associated with better outcomes in HNSCC^29–33^. However, these studies largely included HPV-positive HNSCCs, which is more often associated with greater densities of CD8^+^ T cells and improved survival^22–24^, thus unsurprising that our results differ. However, our results are concordant with a recent study in HNSCC that reported longer survival was associated with patients who had experienced an increase in CD8^+^ tumor-infiltrating lymphocytes from their primary to recurrent tumor^34^. Another study in HNSCC found a similar trend between increased CD8^+^ T cell infiltration, longer survival, presence of specific neoantigens, and increased cytolytic activity in recurrent tumors^35^. Notably, the four patients in our cohort that experienced the greatest decrease in CD8^+^ T cell density in recurrence had TNM stage 4 primary tumors, while patients that experienced an increase in CD8^+^ T cell density in recurrence included TNM stages 1, 2, 3, and 4.

### Quantifying the spatial organization of neoplastic and immune cells

Prior studies reported the abundance of various cell types, including CD8^+^ T cells^33^, CD4^+^ regulatory T cells^36^, and macrophages^37^ to be associated with survival in HNSCC. We analyzed the average density of each cell type across primary tumors for their correlations with PFS, but found no significant association with PFS for any single cell type (p>0.159). Given the prognostic potential of TiME cellular spatial organization as has been reported in other cancer types^5-13^, we quantified the spatial organization of cells within tumor regions and examined the association of the spatial features with clinical outcome.

We first deployed a mixing score, used previously to analyze immune cell spatial compartmentalization in triple negative breast cancers^4^. The mixing score measures the enrichment of neoplastic tumor-immune cell proximity relative to immune-immune cell proximity within a set distance. We quantified the number of immune and neoplastic tumor cells within 15 µm of each other, divided by the number of immune cells within 15 µm from another immune cell. Each region was labeled as mixed or compartmentalized using the median mixing score value for all primary tumors as the threshold (**Figure 3a,b**; see Methods). This threshold classified tumor regions as mixed if at least one neoplastic tumor cell was within 15 µm from an immune cell for approximately every ten immune cells within 15 µm from another immune cell. Tumor regions were considered compartmentalized when this ratio was smaller. Regions with fewer than 250 CD45^+^ immune cells per 800^2^ µm^2^ present were labeled as cold, utilizing the same immune cell density threshold from the original study^4^. Of the 47 total tumor regions, 25 were mixed, 20 were compartmentalized, and two were cold.

**Figure 3.**
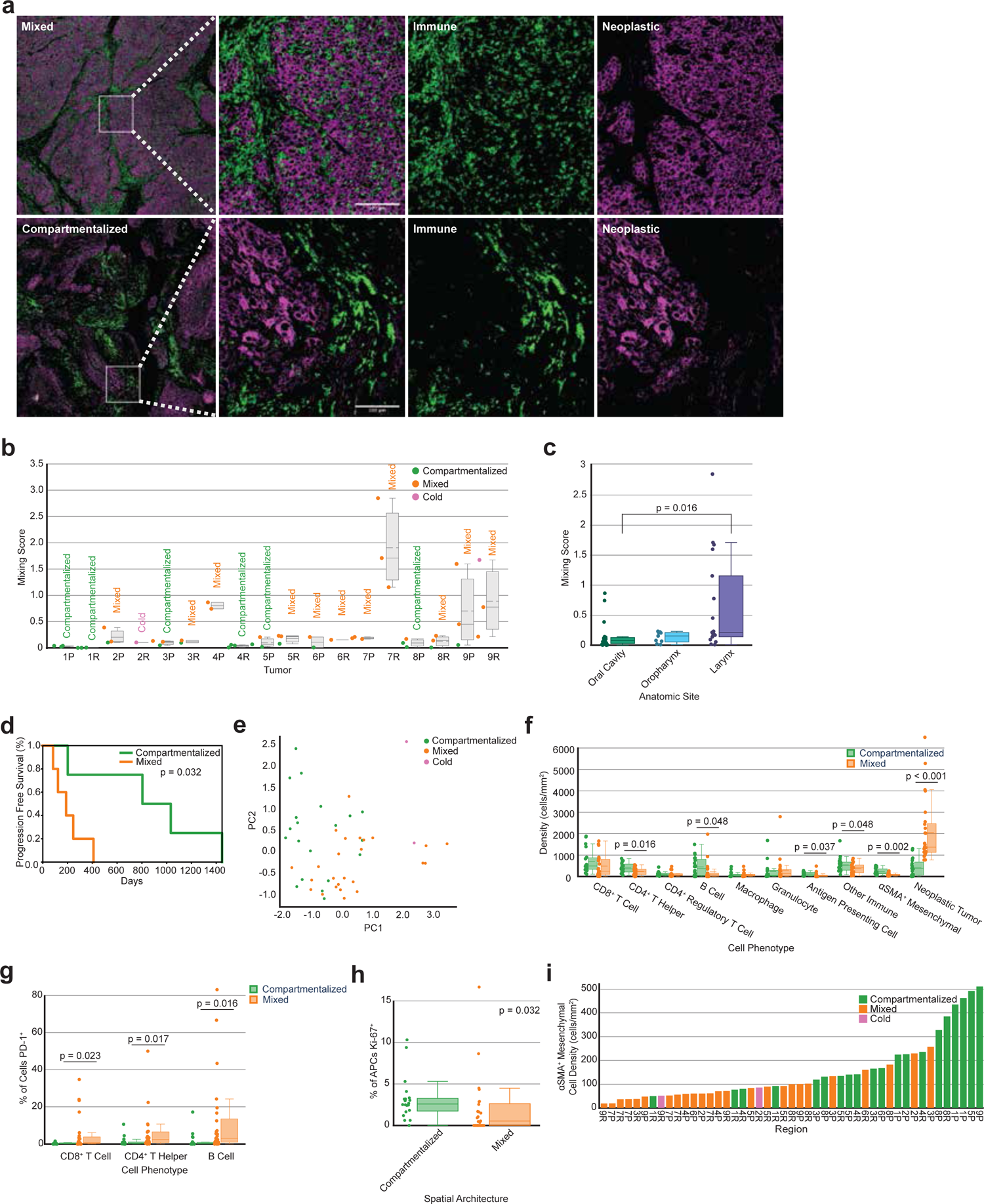
Mixing score quantifies the spatial organization of tumors. **a**, mIHC images of a representative mixed tumor region (top) versus a compartmentalized tumor region (bottom). Leftmost panel shows tumor regions with neoplastic tumor cells (purple) and CD45^+^ immune cells (green). Remaining panels show zoomed in areas of mixing (top) and compartmentalization (bottom), first with both cell populations present and then separated out. **b**, Box plot showing the mixing scores across all primary and recurrent tumors (n=18). Each dot (n=47) represents the mixing score for one tumor region and is colored according to its spatial architecture. The average spatial architecture designation for the overall tumor is printed above each box. **c**, Box plot showing the mixing score of each tumor region split by the anatomic site of its resection. P-value calculated using a one-way ANOVA multi-group significance test followed by a Tukey honestly significant difference post-hoc test. **d**, Kaplan-Meier curve of progression free survival for patients split by the mixing score of their primary tumors. Patients were split on the median value. P-value calculated using the log-rank test. **e**, Principal component analysis on cellular density following a log10+1 transformation. Each dot (n=47) represents one tumor region and is colored according to the region’s spatial architecture. **f**, Box plot showing the density of each cell type split by the tumor region’s spatial architecture. Each dot represents the density of that cell type for one region (n=47 per cell type). Statistical significance calculated using independent one-tailed t-tests for cell types whose differences follow a normal distribution and non-parametric one-tailed t-tests (Mann-Whitney U Test) for cell types whose differences do not follow a normal distribution. P-values were corrected using the Benjamini-Hochberg procedure. **g**, PD-1 expression on CD8^+^ T cells, CD4^+^ T helper cells, and B cells by spatial architecture. Each dot represents the percentage of each cell type positive for PD-1 for a single tumor region (n=45, excluding cold regions). P-values calculated using a one-tailed Mann-Whitney U test and corrected using the Benjamini-Hochberg procedure. **h**, Ki-67 expression on APCs by spatial architecture. Each dot represents the percentage of cells positive for Ki-67 for a single tumor region (n=45, excluding cold regions). P-values calculated using a Mann-Whitney U test and corrected using the Benjamini-Hochberg procedure. **i**, Bar chart showing the density of ⍺SMA^+^ mesenchymal cells present per tumor region. Bars are ordered by ⍺SMA^+^ cell density and are colored according to the region’s spatial architecture.

After calculating the mixing score for each region, we examined the spatial heterogeneity of our cohort. Our tumor compositional heterogeneity analyses revealed that intra-tumoral and intra-patient heterogeneity was less than inter-patient heterogeneity. To determine whether this observation held for spatial organization heterogeneity, we compared each region’s mixing score to five groups of average mixing scores, which were computed from the same five groups as the analysis in Figure 1d: Intra-Tumor (P or R only), Intra-Patient (P and R), Inter-Patient (P or R only), Inter-Patient (Same Anatomic Site), and Inter-Patient (all). Contrary to our analysis in Figure 1d, the mixing score is only one feature, not a distribution of features, thus we used the difference in mixing scores, rather than the KL divergence. In addition, we calculated the absolute values of these differences as a way to normalize the data in order to capture the degree of difference in spatial organization, allowing us to subsequently test for differences across the five levels of heterogeneity. We found there to be less intra-tumoral heterogeneity than inter-patient heterogeneity (**Supplemental Figure 2a**). This result indicates that, in terms of neoplastic tumor-immune cell mixing, tumor regions resemble regions sampled from the same tumor more than regions sampled from tumors of other patients.

We then considered whether tumor regions sampled from the same anatomic site differed in their spatial organization and found that of the three anatomic sites, tumor regions from the oral cavity contained significantly different average mixing scores than tumor regions from the larynx (p=0.016, **Figure 3c**). No significant differences were found in average mixing scores between the oral cavity and the oropharynx or the larynx and the oropharynx. Regions from larynx tumors did exhibit a greater range of mixing scores than oral cavity or oropharynx (**Figure 3c**) indicating that larynx tumors exhibit greater spatial heterogeneity in terms of neoplastic tumor-immune cell proximity. We found no significant difference in the mixing scores of primary versus recurrent tumors (Wilcoxon signed rank test, p=0.441).

### Spatial compartmentalization associated with longer progression free survival

To investigate how spatial mixing correlated with patient outcome, we averaged the mixing scores across regions and assigned a final mixing score and spatial label for each tumor. Patients with more compartmentalization between neoplastic cells and immune cells in their primary tumors exhibited significantly longer PFS than those with greater mixing between these cell types (p=0.032, **Figure 3d**). We then examined how the average mixing score of the tumors related to the TNM stage and anatomic site of each primary tumor. Four of the five mixed primary tumors were TNM stage 4, and one was TNM stage 1. The four compartmentalized primary tumors were TNM stages 1, 2, 3, and 4. All anatomic sites were present in both mixed and compartmentalized spatial architecture groups (mixed: 2 oral cavity, 1 oropharynx, 2 larynx; compartmentalized: 2 oral cavity, 1 oropharynx, 1 larynx), indicating no single anatomic site had predominantly mixed or compartmentalized spatial architecture.

### Spatial architecture associated with cellular composition

We next explored how cellular composition related to spatial organization, in an effort to explain the association found between mixing score and PFS. By coloring the initial PCA on TiME composition by mixing score, we found that tumor regions from the same mixing group clustered together (**Figure 3e**). Due to this association between cellular composition and spatial organization, we wondered whether certain cell types would be more frequent in mixed or compartmentalized tumors. Given how the mixing score is calculated, we hypothesized that compartmentalized tumor regions would have greater densities of immune cells than mixed regions, while mixed tumor regions would have greater densities of neoplastic tumor cells than compartmentalized regions, and found this to be the case (**Figure 3f**). Namely, compartmentalized tumor regions had greater densities of CD4^+^ T helper cells (p=0.016), B cells (p=0.048), antigen presenting cells (APCs) (p=0.037), and other CD45^+^ immune cells (p=0.048) than mixed tumor regions. Given the role of CD4^+^ T helper cells, B cells, and other MHCII^+^ immune cells in antigen presentation, these results could indicate enhanced antigen presentation in compartmentalized tumor regions as compared to mixed regions. Conversely, mixed tumor regions contained greater densities of neoplastic tumor cells than compartmentalized regions (p<0.001).

We further examined associations between functional phenotypes of leukocytes and mixing scores for each tumor region. Mixed regions contained a greater proportion of lymphocytes, including CD8^+^ T cells (p=0.023), CD4^+^ T helper cells (p=0.017), and B cells (p=0.016), expressing the immunoregulatory protein programmed death ligand (PD)-1 than compartmentalized regions (**Figure 3g**). PD-1 is recognized as an indicator of T cell antigen experience, whereas its expression on B cells has been reported to suppress T cell effector function^38, 39^, indicating a suppressive and potentially dysfunctional immune environment in mixed tumors. On the contrary, compartmentalized regions contained a greater proportion of APCs expressing the proliferation marker Ki-67 than mixed regions (p=0.032, **Figure 3h**), supporting the notion that antigen presentation is a key feature of compartmentalized regions. We performed a bootstrapping analysis to confirm the robustness of these results and demonstrate that no one tumor was biasing the functional marker results (**Supplemental Figure 2b,c,d,e**).

In addition to identifying differences between spatial architectures and their respective immune and neoplastic tumor cell densities and functional marker expressions, we found that tumor regions with more compartmentalization between neoplastic cells and immune cells also contained greater densities of αSMA^+^ mesenchymal cells, as compared to those with higher mixing (p=0.002, **Figure 3f**). This result was intriguing because these cells were not included when computing the mixing score, yet there is a clear association between αSMA^+^ cell density and the tumor’s spatial organization (**Figure 3i**).

### αSMA^+^ mesenchymal spatial cellular neighborhoods reveal spatial landscapes associated with progression free survival advantage

Given the relationship between mixing score and αSMA^+^ mesenchymal cell density, we deployed a cellular neighborhood clustering analysis to identify which cell types were spatially proximal to αSMA^+^ cells across tumors in order to gain a better understanding of whether these cells were contributing to the neoplastic tumor-immune spatial compartmentalization observed. This analysis first involved calculating neighborhoods, which were defined as physical groupings of cells within a set distance threshold from a seed cell (**Figure 4a**). Each cell within the distance threshold was deemed a neighbor of the seed cell, contributing to that seed cell’s neighborhood’s composition. After identifying neighborhoods for each seed cell present across all tumor regions, neighborhoods were grouped with K-means clustering, using the fraction of each cell type present in the neighborhoods as the clustering features. This revealed clusters of αSMA^+^ cell neighborhoods with similar cellular makeups across all tumor regions.

**Figure 4.**
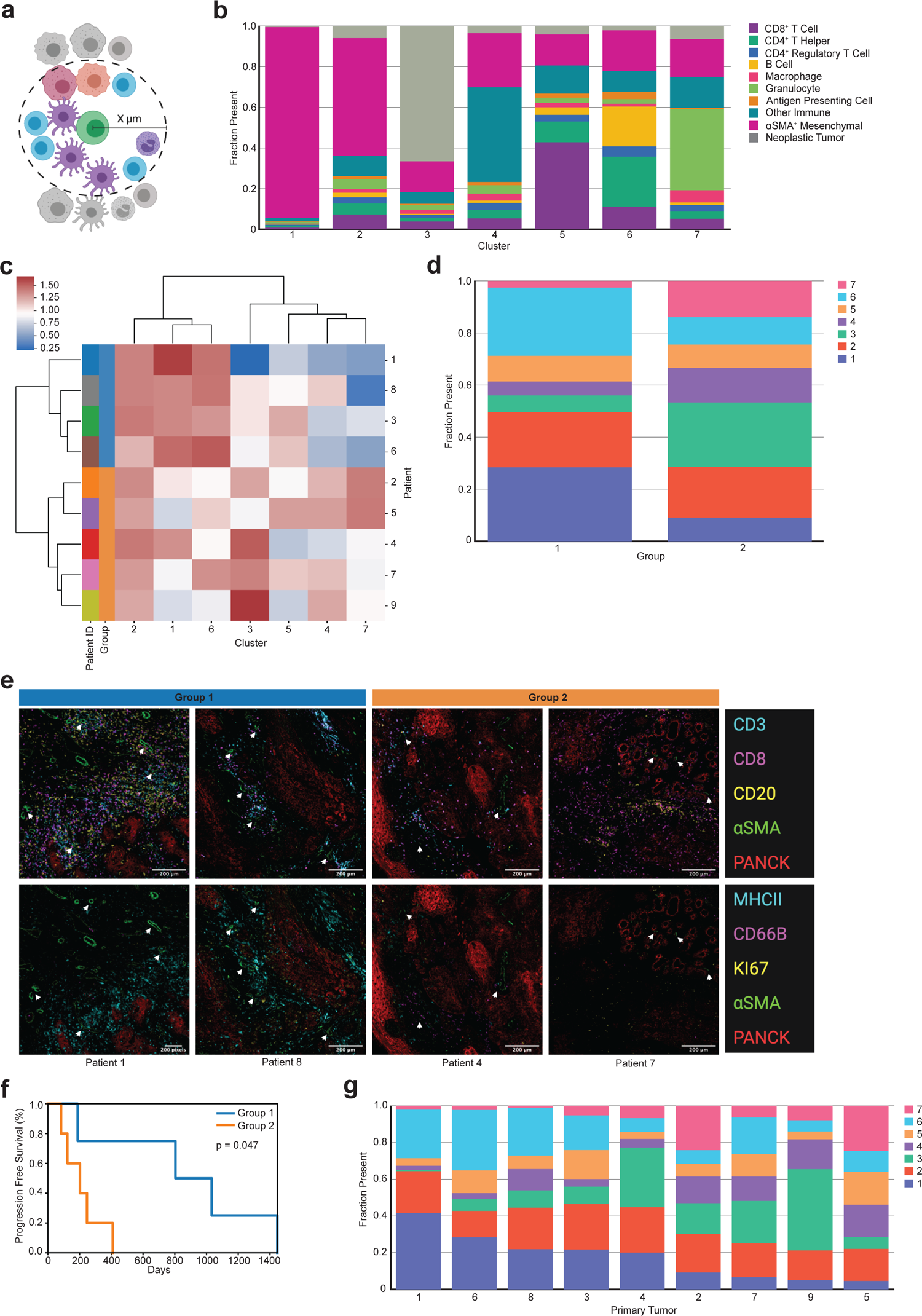
αSMA^+^ mesenchymal cellular neighborhood clustering. **a**, Cellular neighborhoods were defined by drawing a circle of a specified radius around each seed cell (green) of a designated phenotype. Cells whose centers were inside the circle were considered neighbors of that seed cell. This figure was created using BioRender.com. **b**, Stacked bar chart showing the average cellular composition of each αSMA^+^ mesenchymal cell neighborhood cluster (n=7). Bars are colored by cell type and represent the average fraction (out of 1.0) of each cell type present in the neighborhoods belonging to each cluster. **c**, Heatmap of αSMA^+^ cell neighborhood clusters present averaged across primary tumors. Rows are primary tumors that are ordered by the hierarchical clustering of their average of αSMA^+^ neighborhood cluster presence. Columns are the αSMA^+^ cell neighborhood clusters used as clustering features. Percent (out of 100) of αSMA^+^ neighborhood clusters was normalized using a log10+1 transformation before clustering. Leftmost column is color coded by patient. **d**, Stacked bar chart showing the average proportion (out of 1.0) of αSMA^+^ cell neighborhood clusters present in each of the two hierarchically clustered groups of primary tumors. **e**, Panel of mIHC images containing two merged pseudo-colored images of one region from 4 patients visually illustrates representative regions from patients in group 1 and group 2. The top panel of images are merged pseudo-colored stains containing CD3, CD8, CD20, αSMA, and PANCK. The bottom panel of images are the same regions as the top panel with merged pseudo-colored stains containing MHCII, CD66B, Ki-67, αSMA, and PANCK. White arrows point out representative αSMA^+^ cells. **f**, Kaplan-Meier curve of progression free survival for patients grouped together in the hierarchical clustering of their primary tumor αSMA^+^ mesenchymal cell neighborhood abundance. P-value calculated using the log-rank test. **g**, Stacked bar chart showing the average proportion (out of 1.0) of αSMA^+^ cell neighborhood clusters present in each of the primary tumors (n=9). Tumors are ordered by descending proportions of cluster 1.

We applied a neighborhood clustering analysis with αSMA^+^ cells as the seed cells with a distance threshold of 30 µm, as this produced neighborhoods with an average of approximately ten neighbor cells. A recent study involving cellular neighborhood analyses employed a method that selected the ten nearest spatial neighbors of the seed cell, regardless of the distance between the seed cell and its neighbors^7^. By setting a distance threshold of 30 µm, our method required the cells be close, if not directly touching, while still capturing enough neighbors to cluster on. Any αSMA^+^ cell that did not have any neighbors was removed from downstream clustering analyses. Clustering results yielded seven groups, each different in their average composition of αSMA^+^ neighborhoods (**Figure 4b**). Clusters 1 and 2 contained mostly other αSMA^+^ cells comprising the neighborhoods; in fact, cluster 1 was almost exclusively made up of αSMA^+^ cells. Cluster 3 contained the greatest proportion of neoplastic tumor cell neighbors. Clusters 4, 5, 6, and 7 were all comprised of roughly 75% immune cells as neighbors, although they differed in the types of immune cells present. Cluster 4 was defined by a majority of other CD45^+^ immune cells not explicitly defined within our gating strategy. To elucidate marker expression within these other CD45^+^ cells, we performed a post-hoc t-distributed stochastic neighbor embedding (t-SNE) analysis on cells classified as ‘other immune,’ and found them to contain a large population of CD163^+^ cells and a smaller population of mast cells (tryptase^+^) (**Supplemental Figure 3a**). Cluster 5 contained primarily CD8^+^ T cells as the dominant immune population.

Cluster 6 consisted of CD4^+^ T helper cells, B cells, and the greatest proportion of APCs of any cluster. Finally, cluster 7 was defined by its large proportion of granulocytes.

We confirmed that no single tumor region, entire tumor, or patient dominated any of the αSMA^+^ neighborhood clusters by examining the percent contribution of each of the seven clusters per 47 tumor regions, eighteen tumors, and nine patients. While clusters were present in varying degrees across tumor regions, our results indicated that no region (**Supplemental Figure 3b**), tumor (**Supplemental Figure 3c**), or patient (**Supplemental Figure 3d**) was solely responsible for giving rise to any of the clusters. We also confirmed that all seven clusters were present in tumors collected from each of the three anatomic sites (**Supplemental Figure 3e**), as well as each of the four TNM stages (**Supplemental Figure 3f**).

To identify groups of patients with primary tumors of similar αSMA^+^ cellular neighborhoods, we performed unsupervised hierarchical clustering on the normalized average αSMA^+^ cellular neighborhood composition across the nine primary tumors. This resulted in two groups of patients, differing in proportional compositions of αSMA^+^ cellular neighborhoods (**Figure 4c**). On average, both groups had roughly 20% of their αSMA^+^ cells assigned to cluster 2 and roughly 10% of their αSMA^+^ cells assigned to cluster 5 (**Figure 4d**). However, the two groups differed in that group 1 (blue) included patients with αSMA^+^ cells predominantly assigned to clusters 1 and 6, meaning their αSMA^+^ cells were primarily surrounded by CD4^+^ T helper cells, B cells, and other αSMA^+^ cells. On the contrary, group 2 (orange) included patients with more of their αSMA^+^ cells assigned to clusters 3, 4, and 7, meaning their αSMA^+^ cells were mostly surrounded by neoplastic tumor cells, other immune cells, and granulocytes. Consistent with our results, visualization of tissue regions illustrates primary tumors in group 1 containing more αSMA^+^ stromal cells overall, frequently neighboring CD4^+^ T helper cells and B cells, with greater MHCII positivity (**Figure 4e**). Primary tumors in group 2 contained less structured αSMA^+^ stromal cells, fewer neighboring immune cells, and less MHCII positivity, differences that were strikingly apparent in the mIHC stained tissue images (**Figure 4e**). Despite both groups containing nearly equal proportions of CD8^+^ T cells neighboring αSMA^+^ cells, group 1 contained higher densities of CD4^+^ T cells and increased MHCII positivity as compared to group 2 (**Figure 4e**).

To determine if the composition of αSMA^+^ cellular neighborhood groups was correlated with clinical outcome, we performed a survival analysis on the two groups of patients. We found that patients in group 1 had significantly longer PFS than patients in group 2 (p=0.047, **Figure 4f**). Patients in group 1 had primary tumors annotated as TNM stages 1, 2, 3, and 4, while patients in group 2 had four primary tumors annotated as TNM stage 4 and one annotated as TNM stage 1. Tumors from all three anatomic sites were represented in both groups. An analysis of the proportions of αSMA^+^ neighborhood clusters present in each of the nine primary tumors revealed the varying degrees to which each of the seven clusters were present in each of the tumors (**Figure 4g**).

Notably, we found there to be a positive correlation between the presence of clusters 1 and 6 (r = +0.69) as well as clusters 4 and 7 (r = +0.66). We found negative correlations between the presence of clusters 1 and 4 (r = −0.88), clusters 3 and 6 (r = −0.69), and clusters 6 and 7 (r = −0.65). Finally, we found the two groups resulting from hierarchical clustering to be associated with mixing status. Specifically, group 1 consisted of 75% compartmentalized tumors, and group 2 consisted of 80% mixed tumors. Overall, these results describe interesting spatial relationships between immune cells, mesenchymal stroma, and neoplastic tumor cells, indicating increased antigen presentation and immune activity associated with compartmentalization and progression free survival.

## DISCUSSION

The significant role that TiME cellular composition plays in tumor progression and response to therapy has been accepted for over a decade^40^. However, recent findings powered by single-cell proteomics imaging technologies have found that the spatial organization of the cells present in the TiME also plays a critical role^4-13^. TiME spatial quantifications are just beginning to provide novel insights into tumor biology, and thus it is still unclear exactly which spatial features are important in dictating response to therapy or clinical outcome, as well as whether these features are shared across cancer types, and how they could be leveraged for therapeutic decisions and patient stratification. Here, we leveraged single-cell spatial proteomics data generated by our mIHC immunoassay-based imaging platform to quantitatively assess the TiME of nine matched primary and recurrent HPV(-) HNSCCs in order to demonstrate the use of spatial features in disease prognosis. Our results provide novel insights into the heterogeneity and spatial landscape of HPV(-) HNSCCs, and highlight possible TiME spatial landscapes that may impact clinical outcome across cancer types.

We found concordance between the results from our heterogeneity and composition analyses and those from several other studies. Our results are similar to a study that clustered HNSCC biopsies based on their neoplastic and immune gene signatures as determined by RNA-sequencing, and reported that samples from the same patient were more similar to each other than samples from different patients^41^. This trend has also been observed across cancer types, including melanoma^42^, hepatocellular carcinoma^43^, pancreatic ductal adenocarcinoma^44^, and breast cancer^45^. Moreover, we found that patients who experienced an increase in CD8^+^ T cell density from their primary to recurrent tumors were associated with improved PFS, which has previously been reported in head and neck cancer^34, 35^. Conversely, patients experiencing the greatest decrease in CD8^+^ T cell density in recurrence had TNM stage 4 primary tumors. This could indicate that later staged primary tumors are better equipped to evade immune attack in recurrence, and a therapy to elicit an anti-tumor immune response may be beneficial.

Despite the fact that HPV(-) HNSCCs often contain limited immune infiltrates^22^, our analyses revealed significant differences in the immune cell spatial organization within these tumors that were associated with progression free survival, highlighting the importance of considering the spatial context of the TiME. Most strikingly, patients whose primary tumors contained more compartmentalization between their neoplastic tumor cell and immune cell populations demonstrated longer PFS. This correlation was also identified in a similar analysis of triple negative breast cancer patients^4^, indicating that a compartmentalized spatial architecture may play a favorable role in survival across cancer types. Our analyses of TiME composition and functional marker expression indicate there is likely more antigen presentation and less immunoregulation in regions of compartmentalization rather than in regions of mixing. This favorable immune landscape in compartmentalized tumors may contribute to improved survival.

On the contrary, four of the five of the mixed primary tumors were classified as TNM stage 4, which may contribute to shortened survival for these patients. However, given that compartmentalized tumors were not all classified as an early TNM stage and instead ranged from TNM stage 1 to 4, it is difficult to determine the exact association between neoplastic tumor and immune cell mixing versus TNM stage, and further investigation is warranted to understand these correlations and impact on prognosis.

Prior studies have identified a mesenchymal HNSCC subtype^46, 47^; our αSMA^+^ mesenchymal cellular neighborhood analyses highlight the importance of considering how these cells are organized within the TiME, beyond simply considering their presence in the tumor. This could reveal more precise mesenchymal subtypes for improved stratification for patient care. αSMA is a common marker for cancer-associated fibroblasts (CAFs)^20, 48^, whose presence in tumors, including HNSCC, tends to be associated with tumor progression, metastasis, and resistance to therapy^49–54^.

However, despite compartmentalized tumors having increased αSMA^+^ cell density, patients with compartmentalized primary tumors demonstrated longer PFS, which indicates that the spatial organization of αSMA^+^ cells may be related to their function.

We found αSMA^+^ cells neighboring immune cells in both mixed and compartmentalized primary tumors, but the types of immune cells near αSMA^+^ cells differed. This is interesting, as emerging research has found that CAFs, which are often defined by their expression of αSMA, can modulate immune cell function within the TiME. With our mIHC platform we were able to identify the differences in the types of immune cells neighboring αSMA^+^ cells and relate these differences to survival, supporting recent research on the impact of CAFs on various immune cell populations. A recent study in melanoma found CAFs to be instrumental in aiding tertiary lymphoid structure (TLS) development^55^; TLS are defined by their large concentration of B cells surrounded by T cells, and their presence in tumors is associated with improved patient outcome^3,56, 57^. In compartmentalized tumors, we noticed the αSMA^+^ cells to be more structured and located neighboring dense pockets of T cells and B cells, perhaps indicating the formation of TLS. Although this antibody panels did not include a biomarker for high endothelial venules to confirm presence of TLS, our results support the notion that CAF-lymphocyte interactions can be clinically beneficial.

Conversely, patients with predominantly mixed primary tumors and shorter PFS contained αSMA^+^ cells primarily located near granulocytes and other CD45^+^ immune cells, many of which were likely CD163^+^ myelomonocytic cells or mast cells. CD163 is a marker for scavenger receptor activity and is commonly used to demark pro-tumor type tumor-associated macrophages and monocytes. Supporting these findings, a study in oral squamous cell carcinoma found the presence of CAFs to be associated with increased presence of CD163^+^ macrophages and worse survival^58^. Another study reported CAFs to be correlated with an increase in monocyte expression of CD163, which in turn suppressed T cell proliferation and increased neoplastic cell proliferation in breast cancer^59^. Mast cells mediate innate and acquired immune response as a part of the myeloid lineage. They have been reported to facilitate neo-vascularization and tumor dissemination in HNSCC, and found to be correlated with increased angiogenesis in advanced HNSCC^60^. However, interactions between mast cells and CAFs is largely unknown; our results indicate this interaction may be associated with increased neoplastic density and worse PFS. Of note, a recent study in melanoma and pancreatic adenocarcinoma found neutrophils, a subclass of granulocytes, to exert pro-tumor effects when in the presence of CAFs^61^, and another study found the combined presence of neutrophils and CAFs to be associated with shortened survival in gastric adenocarcinoma^62^.

Finally, patients with predominantly mixed primary tumors and shorter PFS were found to contain fewer overall stromal cells and greater proportions of those αSMA^+^ cells neighboring neoplastic tumor cells. It has been reported that neoplastic tumor cells in direct contact with CAFs move along tracks laid by the CAFs in the extracellular matrix, promoting tumor growth^63, 64^. It is possible that αSMA^+^ cells in mixed tumors may provide avenues for neoplastic tumor cells to transit, thus leading to more advanced tumor progression. Overall, spatial analyses herein deepened our understanding of neoplastic tumor and immune cell organization relative to each other, and how this organization is related to αSMA^+^ cells working in tandem with many cells, beyond single cell-cell interactions, to impact TiME organization, and ultimately, clinical outcome. The conclusions from our spatial analyses and their relationship to clinical variables are summarized in **Figure 5**.

**Figure 5.**
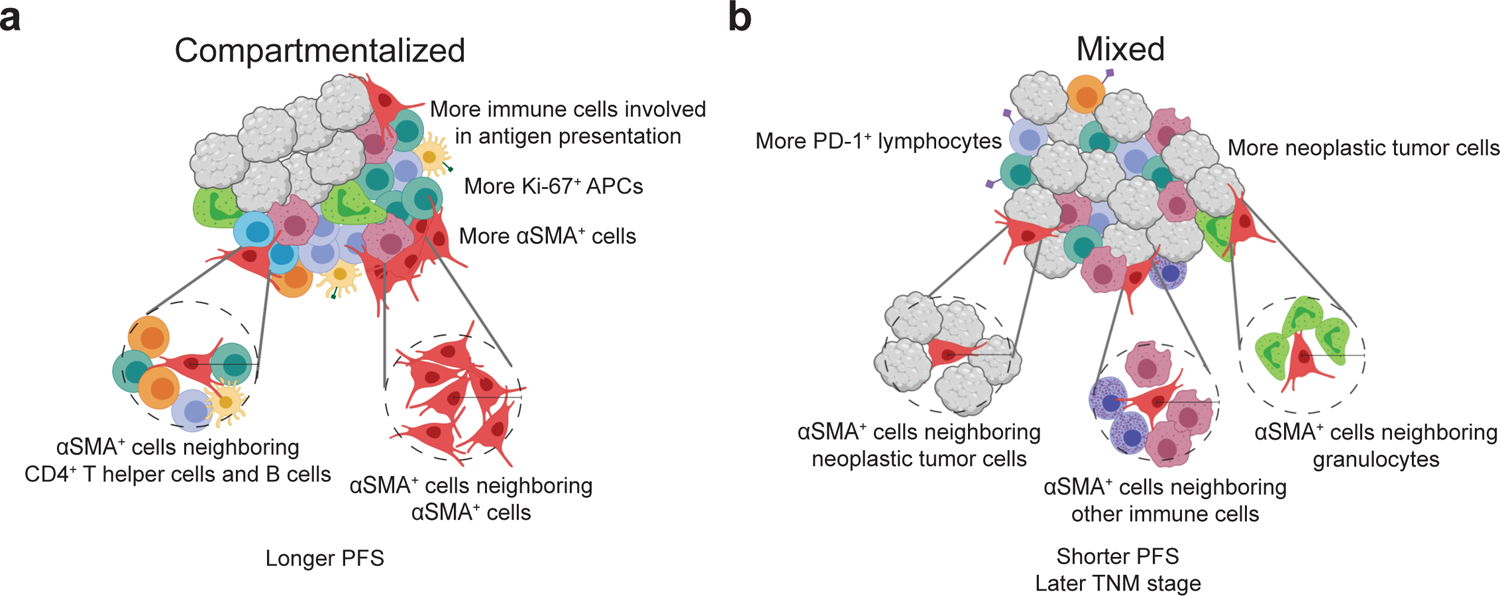
Proposed model of primary HPV(-) HNSCC tumor-immune microenvironments. **a**, Depiction of a tumor with a compartmentalized spatial architecture. These tumors have decreased mixing between immune cells and neoplastic tumor cells and tend to contain greater immune cell density, specifically those involved in antigen presentation, as well as increased density of αSMA^+^ mesenchymal cells. The αSMA^+^ cells present tend to be neighbored by CD4^+^ T helper cells and B cells, as well as other αSMA^+^ cells. Compartmentalized primary tumors were found to be associated with longer progression free survival. **b**, Depiction of a tumor with a mixed spatial architecture. These tumors contain increased mixing between immune cells and neoplastic tumor cells and tend to contain increased neoplastic tumor cells, as well as increased PD-1-positive lymphocytes. The αSMA^+^ cells present tend to be neighbored by neoplastic tumor cells, other immune cells, many of which are CD163^+^ or are mast cells, and granulocytes. Mixed primary tumors were found to be associated with shorter progression free survival and later TNM stage. This figure was created using BioRender.com.

The concordance found among results herein and those of the aforementioned studies provides orthogonal support for our conclusions and indicates that single-cell heterogeneity and spatial organization of tumors may share similarities across different types of cancer and across different molecular assays. Our spatial analyses demonstrate the use of various algorithms to quantify the spatial landscape of tumors using single-cell imaging data. Further, our computational methods provide a framework for future single-cell imaging analyses, as they are applicable to any multiplex imaging assay, including mIHC, co-detection by indexing technology (CODEX)^7^, cyclic immunofluorescence^18, 65, 66^, and imaging mass cytometry^6^. Our results highlight several spatial architectures that may help guide precision medicine approaches for HPV(-) HNSCC patients, including architectures that may help stratify patients who may have shorter PFS, and thus warrant more aggressive therapy or clinical follow-ups. While our results provide evidence that the spatial organization of HPV(-) HNSCC tumors correlates with clinical outcome, future studies with larger cohorts will be needed to evaluate the strength and validity of our observations. Studies with greater representation of anatomic site and stage are also needed to assess the prognostic value of the spatial features identified in this HNSCC cohort. Despite these limitations, this study demonstrates practical analysis strategies that elucidate spatial architecture features for potential use in precision medicine.

## METHODS

### Multiplex immunohistochemistry data generation

mIHC is an immunohistochemical-based imaging platform that evaluates sequentially stained immune lineage epitope-specific antibodies for immunodetection on FFPE tissue sections^16, 17^. Images were stained and processed as described in our previous report^17^. A table of antibodies, species, vendor, and concentration used in staining are previously reported in Table 1 of Banik et al^17^. Briefly, sequentially stained images were co-registered in MATLAB. AEC signal from each antibody stain was extracted and normalized, and the mean intensity of each single cell for each marker was quantified in Cell Profiler. Watershed based nuclei segmentation on hematoxylin staining was used to identify single cells in FIJI. Using a hierarchical gating strategy, single cells were phenotyped using image gating cytometry in FCS Express 7 Image Cytometry RUO (**Supplemental Figure 1a**). A threshold was set on the scatterplot of mean intensity for each marker within the gating strategy, validated by visual live rendering of masked cell objects within the selected gate on extracted marker signals. Cartesian coordinates of each phenotyped cell were maintained relative to the tissue region.

We applied the mIHC pipeline to analyze matched primary and recurrent FFPE tissue specimens from nine HPV(-) HNSCC patients. Each patient underwent surgical resection of their primary tumor prior to beginning a regimen of chemotherapy and radiation therapy. Upon recurrence, the patient underwent another surgical resection of their recurrent tumor. Tissue specimens for each patient were obtained from the Oregon Health & Science University Knight Biolibrary and were deidentified and coded with a unique identifier prior to analysis. Patient demographic and clinical data including HPV status, tobacco and alcohol use, treatment regimens, and survival outcomes were collected. All HNSCC tumors were staged according to the 8th edition AJCC/UIC TNM classification and cohort characteristics are shown in **Table 1** as reported in Banik et al^17^. All studies involving human tissue were approved by institutional IRB (protocol #809 and #3609), and written informed consent was obtained.

### Tumor heterogeneity analyses

The Kullback-Leibler divergence was calculated using the entropy function from the Scipy Python package^67^. The distribution of each individual tumor region’s cellular composition was compared to the average of five different cellular composition distributions: the average distribution for the region’s tumor, the average distribution for the patient’s primary and recurrent tumors, the average distribution across all tumors of the same timepoint in the cohort (primary or recurrent), the average distribution across all tumors resected from the same anatomic site in the cohort, and the average distribution across all tumors from all patients in the cohort (primary and recurrent combined). Log base 2 was used for the calculation. Unsupervised hierarchical clustering of each tumor region was performed using the log10+1 normalized density of each cell type present as the features. Euclidean distance was used to determine distances between observations, and the Ward method was used for the linkage.

### TiME compositional change clustering analysis

Unsupervised hierarchical clustering of each patient was performed using the normalized change in density of each cell type as the features. Euclidean distance was used to determine distances between observations, and the Ward method was used for the linkage. Normalization in the change in density was computing by first calculating the absolute value of the raw change in density for each cell type. These values were then normalized to a range of [0,1]. Finally, the values that were originally negative (decreasing change), were flipped to be negative values again, such that all values ranged from [-1,1] with zero representing no change.

### Mixing score analysis

Keren et al. developed a mixing score to quantify the ratio of neoplastic and immune cell spatial interactions^4^. This score is defined as the number of interactions between neoplastic tumor cells and immune cells divided by the number of interactions between immune cells and another immune cell within a tumor region. We defined there to be an interaction between two cells if their centers [the (x,y) coordinates provided by mIHC] were within 15 µm from one another. We used the median mixing score value (0.107) for all primary tumors as the threshold to distinguish between mixed and compartmentalized spatial organization groups. Tumor regions with a mixing score of greater than 0.107 were defined as mixed. Tumor regions with a mixing score of less than 0.107 were defined as compartmentalized. We used a density threshold of less than 250 immune cells per 800^2^ µm^2^ to define tumor regions as cold. We chose this threshold to match that used by Keren et al.

### Functional marker bootstrapping analyses

Bootstrapping analyses involved the following steps. First, one tumor region per eighteen tumor samples was randomly selected. The regions were then split into two groups based on their mixed or compartmentalized spatial architecture designation, and the average proportion of the specified cell population expressing the specified functional marker was calculated for each group. Finally, this process was repeated 100 times, yielding 100 values representing the proportion of the specified cell population expressing the specified functional marker for mixed tumor regions and 100 values representing the proportion of the specified cell population expressing the specified functional marker for compartmentalized tumor regions. These values were then compared between the mixed and compartmentalized groups for differences.

### Cellular neighborhood clustering analyses

Cellular neighborhoods were defined by drawing a circle of a specified radius around all seed cells of a given phenotype. Cells whose centers were inside the circle were considered neighbors of that seed cell and contributed to that cell’s neighborhood. All neighborhoods across all tumor regions were then clustered using scikit-learn’s MiniBatchKMeans function^68^ to perform K-means clustering according to their normalized cellular composition. The elbow method was used to determine the number of clusters to form. Each resulting cluster was comprised of neighborhoods with a similar cellular makeup. Unsupervised hierarchical clustering of each primary tumor was performed using the log10+1 normalized proportion (out of 100) of αSMA^+^ cell neighborhood clusters present in the tumor as the features. Euclidean distance was used to determine distances between observations, and the Ward method was used for the linkage.

### Statistics

Independent t-tests were used to determine statistically significant differences for independent samples whose differences followed a normal distribution. Mann-Whitney U tests were used to determine statistically significant differences for independent samples whose differences did not follow a normal distribution. Paired t-tests were used to determine statistically significant differences for paired samples whose differences followed a normal distribution. Wilcoxon signed rank tests were used to determine statistically significant differences for paired samples whose differences did not follow a normal distribution. One-way ANOVA tests were used to determine statistically significant differences for multi-group comparisons. If the ANOVA result was significant, a Tukey honestly significant difference post-hoc test was conducted to determine which groups were significantly different from one another. A Benjamini-Hochberg correction was used to account for multiple hypothesis testing in analyses that involved systematically testing multiple variables. P-values less than 0.05 were considered statistically significant. All statistical calculations were performed with the Scipy and statsmodels packages using Python software^67, 69^.

### Survival analyses

Kaplan-Meier curves were generated and a log-rank test was performed using the lifelines package with Python software^70^. P-values less than 0.05 were considered statistically significant.

### Data and software availability

All of the data produced by our mIHC computational image processing pipeline, including protein abundance, cell phenotype, and cell location information saved in the form of a matrix, in addition to survival data, is available for download at Zenodo (https://doi.org/10.5281/zenodo.5540356). All computational analyses in this study were performed using Python software, version 3.6.5. The code created to produce the results of this study is available at https://github.com/kblise/HNSCC_mIHC_paper.

## ACKNOWLEDGEMENTS

The authors thank Dr. Courtney Betts, Justin Tibbitts, Teresa Beechwood, and Meghan Lavoie for regulatory and technical assistance. The authors thank the OHSU Knight Biostatistics Shared Resource for guidance with statistical calculations. The authors also acknowledge and thank the patients who donated tissue samples for this study.

## Funding

KEB acknowledges funding from the National Cancer Institute (NCI) of the National Institutes of Health under award number 5T32CA254888. LMC acknowledges funding from the NCI awards U01CA224012, R01CA223150, R01CA226909, and R21HD099367 and funding from the Brenden-Colson Center for Pancreatic Care at Oregon Health & Science University (OHSU). JG acknowledges funding under award number U24CA231877. LMC and JG acknowledge funding from the OHSU Knight Cancer Institute, NCI award number U2CCA233280, and the Prospect Creek Foundation to the OHSU SMMART (Serial Measurement of Molecular and Architectural Responses to Therapy) Program. The content is solely the responsibility of the authors and does not necessarily represent the official views of the National Institutes of Health, OHSU, or the Knight Cancer Institute.

## Conflict of Interests

L.M. Coussens is a paid consultant for Cell Signaling Technologies, AbbVie Inc., and Shasqi Inc., received reagent and/or research support from Plexxikon Inc., Pharmacyclics, Inc., Acerta Pharma, LLC, Deciphera Pharmaceuticals, LLC, Genentech, Inc., Roche Glycart AG, Syndax Pharmaceuticals Inc., Innate Pharma, NanoString Technologies, and Cell Signaling Technologies, is a member of the Scientific Advisory Boards of Syndax Pharmaceuticals, Carisma Therapeutics, Zymeworks, Inc, Verseau Therapeutics, Cytomix Therapeutics, Inc., Hibercell., Inc., Alkermes, Inc., and Kineta Inc, and is a member of the Lustgarten Therapeutics Advisory working group. No potential conflicts of interest were disclosed by the other authors.

---

**Supplemental Figure 1a.**
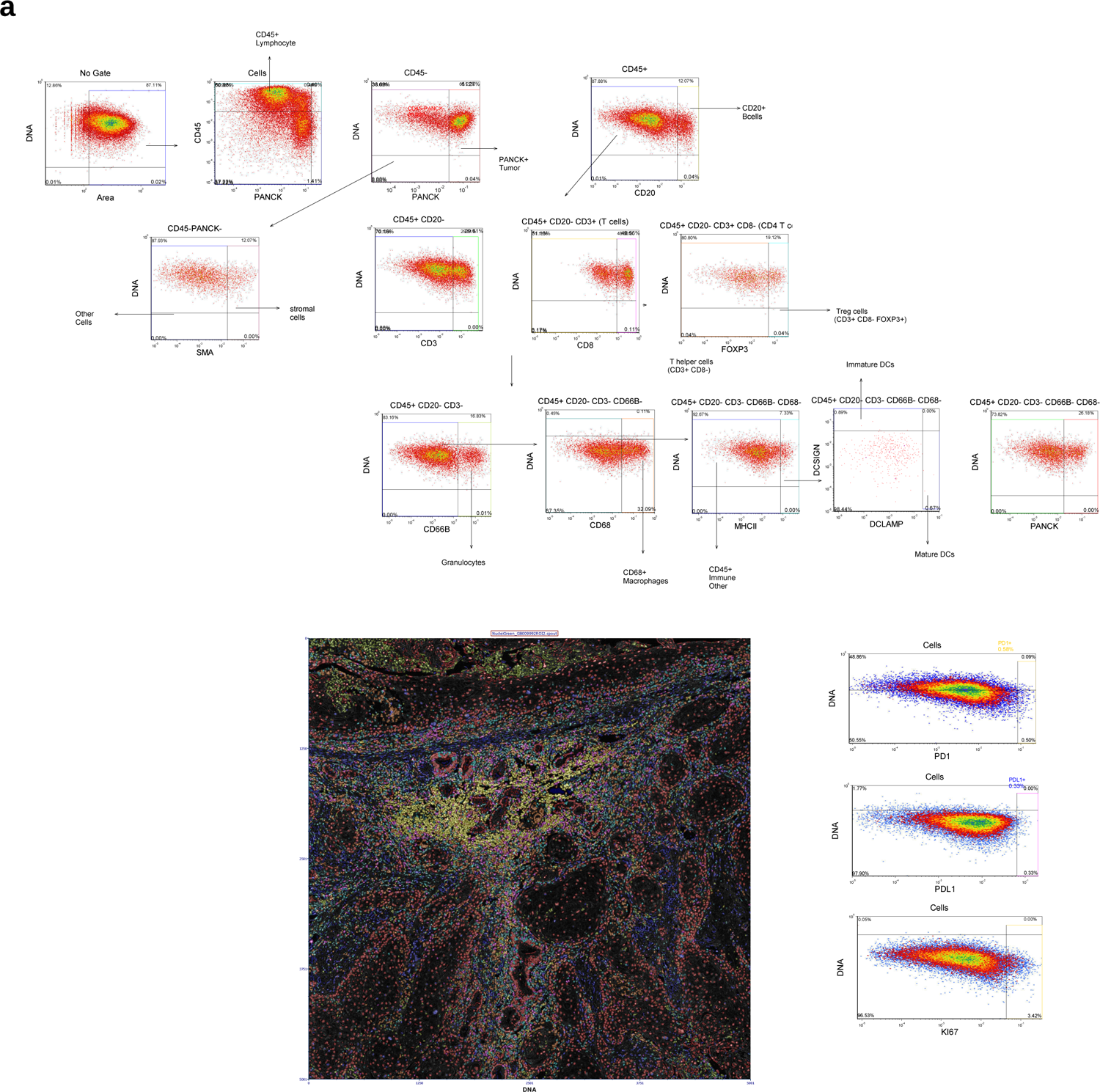
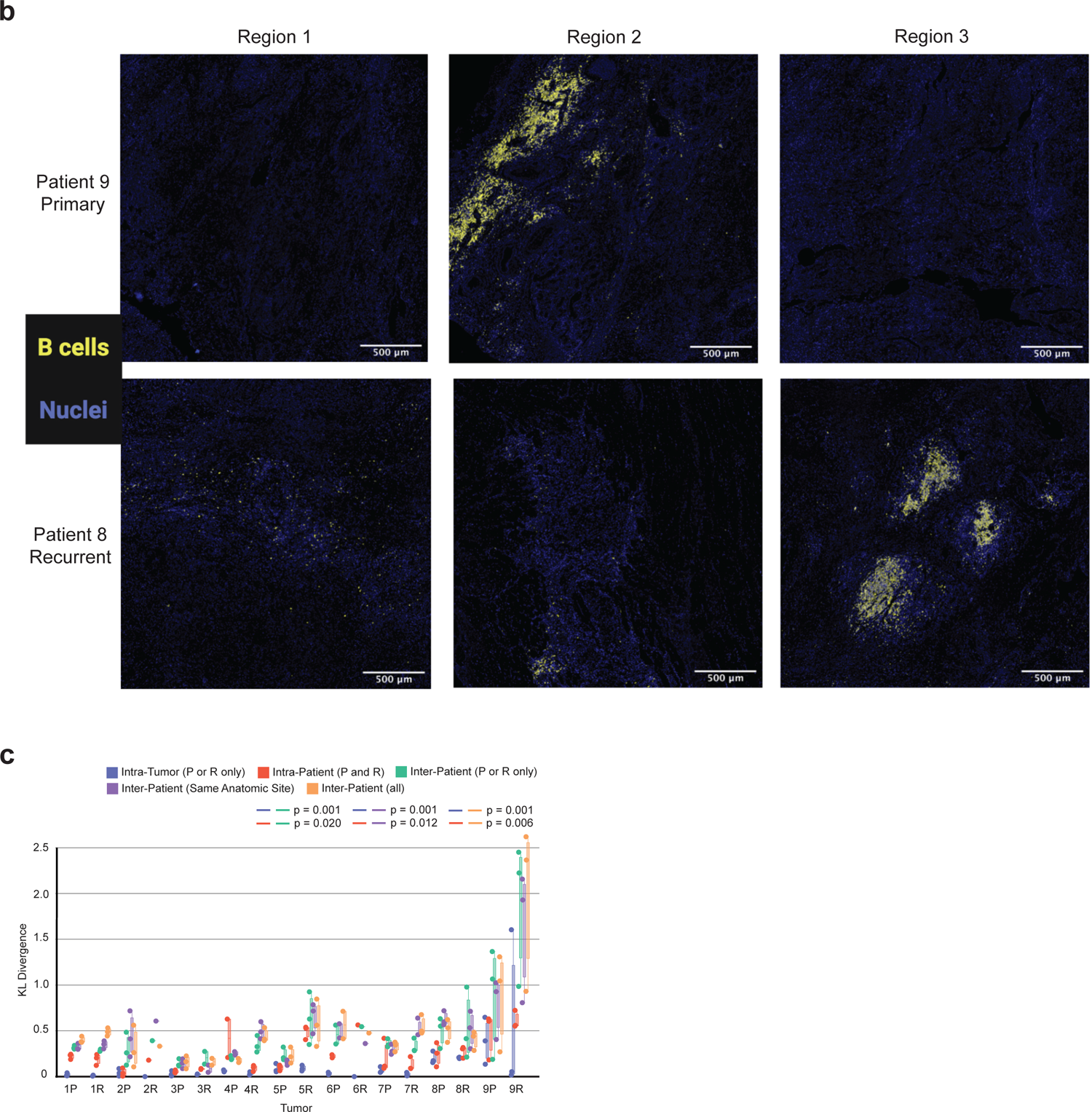
Gating template used to determine cell phenotypes and classification of ten populations using image gating cytometry in FCS Image Cytometry 7 RUO. Scatter plots show single cell mean intensity distributions rescaled from 0-1 on a log10 scale with gated populations in colored boxes. Image plot shows a semi-transparent pseudo-colored mask of each cell object colored by the classified population overlaid on signal extracted nuclei image. The markers used for identification of cell phenotypes are shown in Table 1. Supplemental Figure 1b. mIHC images of six tumor regions selected from two different tumor resections (patient 9 primary and patient 8 recurrent). Blue pseudo color indicates all cell nuclei present in the region (Hematoxylin^+^) and yellow indicates B cells. In both tumors, two regions have low densities of B cells, whereas one region has a high density and spatial concentration of B cells, despite the three regions being sampled from the same tumor. Supplemental Figure 1c. Box plot of the Kullback-Leibler divergences from a single tumor region’s immune cell distribution compared to: the tumor’s average immune cell distribution [Intra-Tumor (P or R only)], the patient’s average immune cell distribution [Intra-Patient (P and R)], the cohort’s average immune cell distribution across tumors of the same timepoint [Inter-Patient (P or R only)], the cohort’s average immune cell distribution across tumors of the same anatomic site [Inter-Patient (Same Anatomic Site)], the cohort’s average immune cell distribution across all tumors from all patients [Inter-Patient (all)]. P-values calculated using a one-way ANOVA multi-group significance test followed by a Tukey honestly significant difference post-hoc test.

**Supplemental Figure 2a.**
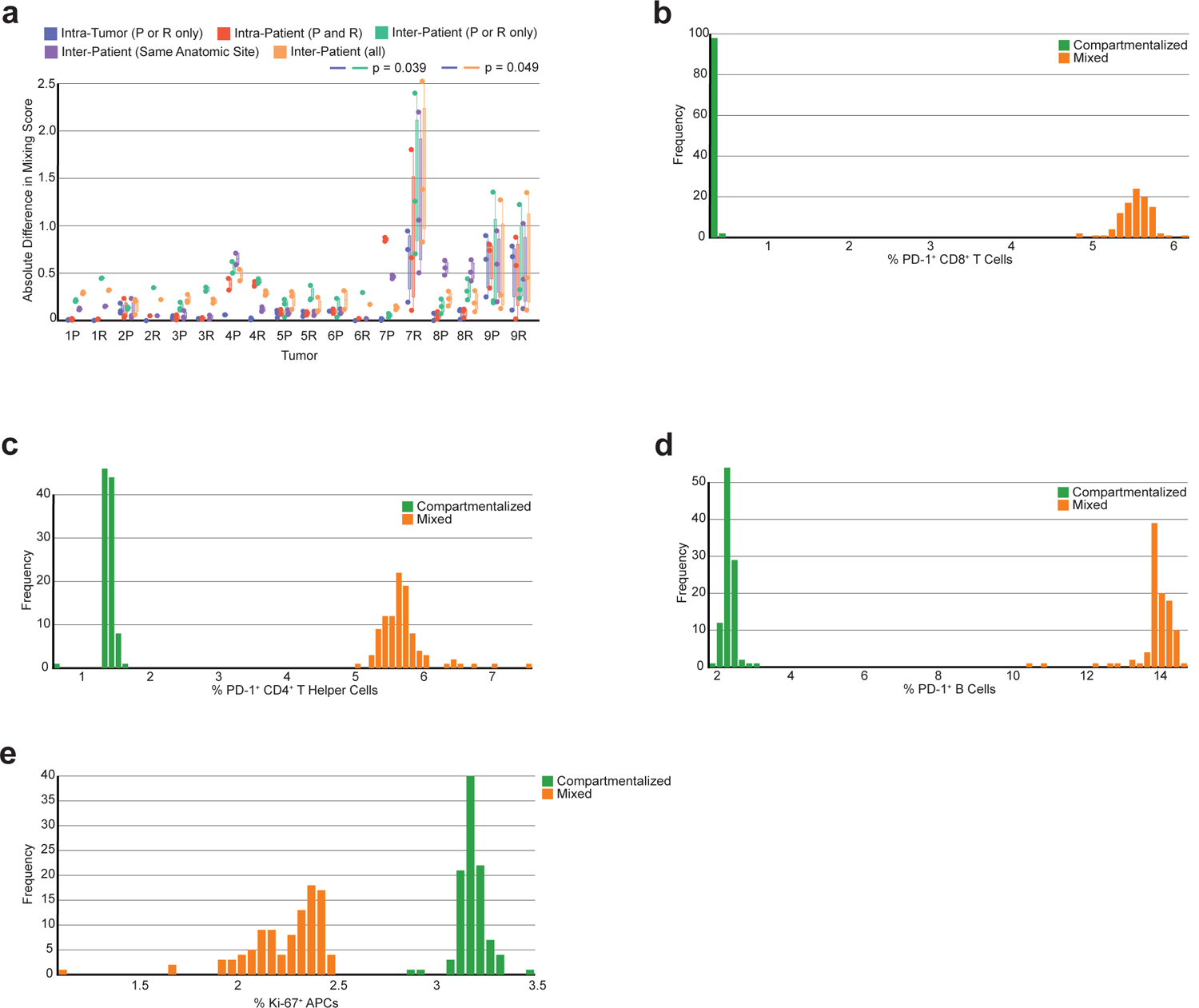
Box plot of the absolute difference from a single tumor region’s mixing score compared to: the tumor’s average mixing score [Intra-Tumor (P or R only)], the patient’s average mixing score [Intra-Patient (P and R)], the cohort’s average mixing score across tumors of the same timepoint [Inter-Patient (P or R only)], the cohort’s average mixing score across tumors of the same anatomic site [Inter-Patient (Same Anatomic Site)], the cohort’s average mixing score across all tumors from all patients [Inter-Patient (all)]. P-value calculated using a one-way ANOVA multi-group significance test. Supplemental Figure 2b. Histogram showing the results of a bootstrapping analysis on the percentage of CD8^+^ T cells expressing PD-1. Bars are colored by spatial architecture. Supplemental Figure 2c. Histogram showing the results of a bootstrapping analysis on the percentage of CD4^+^ T helper cells expressing PD-1. Bars are colored by spatial architecture. Supplemental Figure 2d. Histogram showing the results of a bootstrapping analysis on the percentage of B cells expressing PD-1. Bars are colored by spatial architecture. Supplemental Figure 2e. Histogram showing the results of a bootstrapping analysis on the percentage of APCs expressing Ki-67. Bars are colored by spatial architecture.

**Supplemental Figure 3a.**
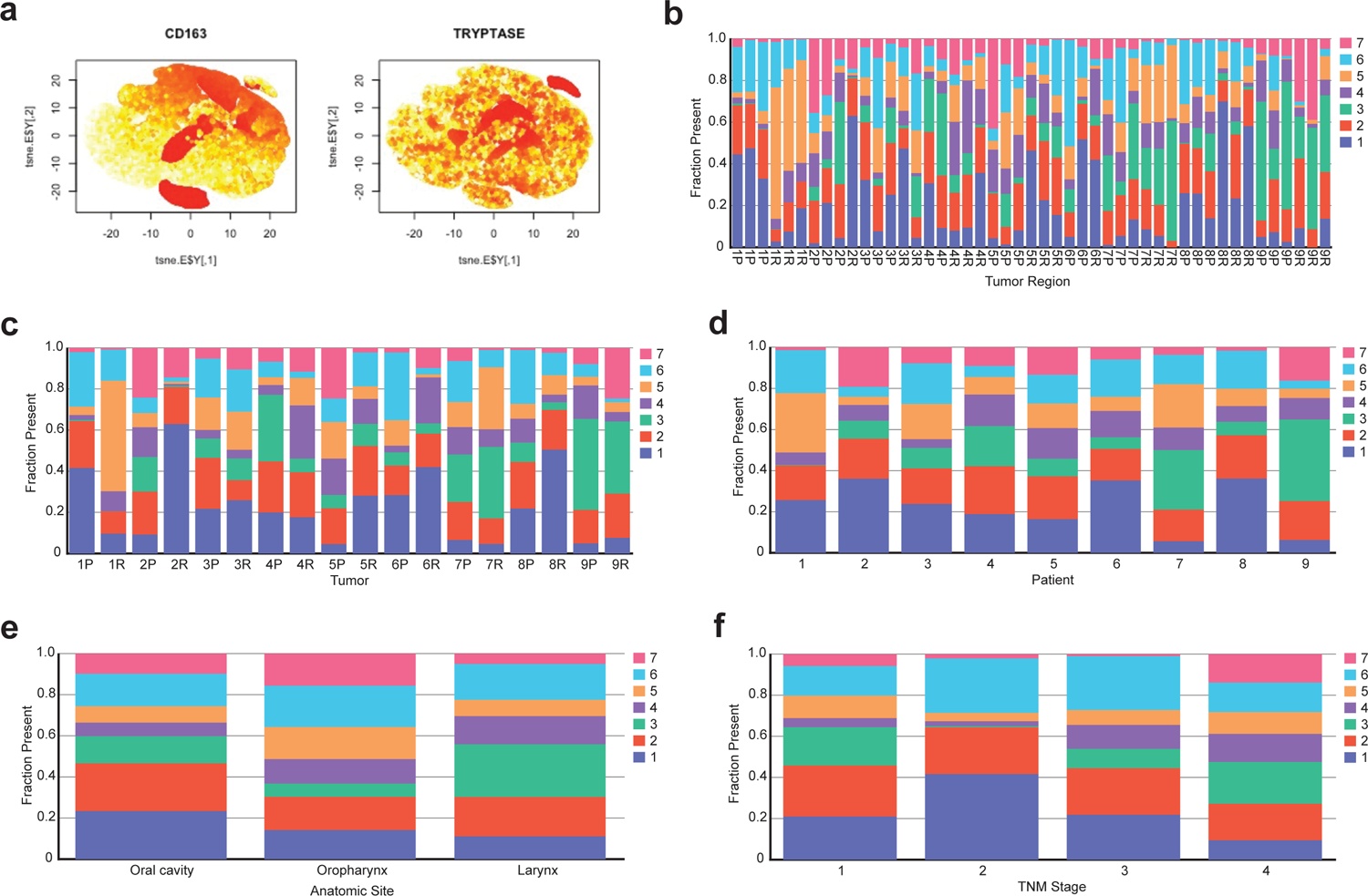
t-SNE representation of single cells from the ‘Other Immune’ class colored by heatmap of mean intensity for CD163 and tryptase (mast cells) show distinct populations of cells expressing high levels of CD163 and tryptase, indicating these cell types are likely present in the ‘Other Immune’ class. Supplemental Figure 3b. Stacked bar chart showing the proportion (out of 1.0) of αSMA^+^ cell neighborhood clusters present in each tumor region (n=47). Supplemental Figure 3c. Stacked bar chart showing the average proportion (out of 1.0) of αSMA^+^ cell neighborhood clusters present in each tumor (n=18). Supplemental Figure 3d. Stacked bar chart showing the proportion (out of 1.0) of αSMA^+^ cell neighborhood clusters present in each patient’s primary and recurrent tumors averaged (n=9). Supplemental Figure 3e. Stacked bar chart showing the average proportion (out of 1.0) of αSMA^+^ cell neighborhood clusters present in tumors collected from the three anatomic sites. Supplemental Figure 3f. Stacked bar chart showing the average proportion (out of 1.0) of αSMA^+^ cell neighborhood clusters present in the primary tumors by their TNM stage.

## Notes

https://doi.org/10.5281/zenodo.5540356

https://github.com/kblise/HNSCC_mIHC_paper

